# The role of African buffalo in the epidemiology of foot-and-mouth disease in sympatric cattle and buffalo populations in Kenya

**DOI:** 10.1101/484808

**Authors:** George Omondi, Francis Gakuya, Jonathan Arzt, Abraham Sangula, Ethan Hartwig, Steven Pauszek, George Smoliga, Barbara Brito, Andres Perez, Vincent Obanda, Kimberly VanderWaal

## Abstract

Transmission of pathogens at wildlife-livestock interfaces poses a substantial challenge to the control of infectious diseases, including for foot-and-mouth disease virus (FMDV) in African buffalo and cattle. The extent to which buffalo play a role in the epidemiology of this virus in livestock populations remains unresolved in East Africa. Here, we show that FMDV occurs at high seroprevalence (~77%) in Kenyan buffalo. In addition, we recovered 80 FMDV VP1 sequences from buffalo, all of which were serotype SAT1 and SAT2, and seventeen FMDV VP1 sequences from cattle, which included serotypes A, O, SAT1 and SAT2. Notably, six individual buffalo were co-infected with both SAT1 and SAT2 serotypes. Our results suggest that transmission of FMDV between sympatric cattle and buffalo is rare. However, viruses from FMDV outbreaks in cattle elsewhere in Kenya were caused by viruses closely related to SAT1 and SAT2 viruses found in buffalo. We also show that the circulation of FMDV in buffalo is influenced by fine-scale geographic features, such as rivers, and that social segregation amongst sympatric herds may limit between-herd transmission. Our results significantly advance knowledge of the ecology and molecular epidemiology of FMDV at wildlife-livestock interfaces in Eastern Africa, and will help to inform the design of control and surveillance strategies for this disease in the region.

## Introduction

Foot-and-mouth disease virus (FMDV) is a vesicular disease of wild and domestic ungulates of the genus *Apthovirus* virus of the family *Picornaviridae*. It is highly transmissible and amongst the most economically important livestock diseases globally. Endemicity of FMDV in East Africa is sustained by inadequate disease control and presence of a wildlife reservoir as a presumed source of infection to livestock (Tekleghiorghis et al., 2016). Understanding its epidemiology in the region is critical to implement effective control strategies. Diseases at wildlife-livestock interfaces in East Africa are a substantial concern for livestock holders and conservationists, and misconceptions arise as to whether wild mammals are the main source of pathogens in domestic stock (Kock, 2005, Caron et al., 2003). In Kenya, rangelands account for 80% of the landmass and are shared by diverse wildlife communities coexisting with pastoralist livestock populations, which is considered a largely beneficial interaction with ecological and trophic benefits (Schieltz and Rubenstein, 2016, Marchant and Lane, 2014). Human populations residing in rangelands depend on livestock production, which is characterized by utilization of the ecosystem through livestock mobility, and shared pasture and water resources. However, high contact rates between domestic and wild animals has implications for interspecies disease sharing and emergence (Kock, 2005).

FMDV can be divided into seven serotypes (O, A, C, SAT 1, SAT 2, SAT3, and Asia 1)(Kitching et al., 1989), with no cross-immunity between serotypes (Kitching et al., 1989). In Kenya, FMDV was first described in 1952, with serotypes O, A, SAT 1 and SAT 2 presently circulating in the country (Wekesa et al., 2015b) and generally in East Africa (FAO., 2016, Sangula et al., 2011). A Bayesian phylogenetic analysis of SAT1 serotypes from Kenya revealed two independent introductions from southern Africa (Sangula et al., 2010); with the Kenya-Tanzania border showing greater transboundary spread of the virus than Kenya-Uganda (Sangula et al., 2010), possibly as a result of pastoralism and seasonal movement of both wild and domestic animals between Kenya and Tanzania. In livestock, FMDV pathogenesis includes an acute symptomatic phase characterized by pyrexia and lesions (starting as vesicles) on the buccal mucus membranes, tongue, teat and feet (Arzt et al., 2011). In some individuals, the virus may persist in the pharyngeal region and potentially serve as a source of infection (Arzt et al., 2011). This persistent infection, also known as the carrier stage, has been shown to last up to 2.5 years in cattle (Salt, 1993, Alexandersen et al., 2002), five months in sheep (Burrows, 1968), and five years in African buffalo (specifically for SAT serotypes)(Condy et al., 1985). The carrier state presupposes an evolutionary host-pathogen interaction allowing for the maintenance of the virus in nature, and has implications for the epidemiology of FMDV in both livestock and wildlife species.

Buffalo are reservoirs for serotypes SAT1, 2 and 3 (Condy et al., 1985) and seroprevalence in East African buffalo is nearly 70% (BronsVoort et al., 2008), but infection causes mild or no clinical signs. It is clear that buffalo are maintenance hosts for SAT serotypes of FMDV, and proximity to wildlife areas is a risk factor for clinical outbreaks in cattle in East Africa (Ayebazibwe et al., 2010a). However, numerous and widespread outbreaks of FMDV in cattle throughout the region suggest that buffalo are not a common source of infection for cattle (Kock et al., 2014, Casey-Bryars et al., 2018). The circumstances under which buffalo-cattle FMDV transmission occurs remain poorly understood (Bastos et al., 2000a, Hargreaves et al., 2004, Ferguson et al., 2013), particularly in East Africa. Although the importance of a buffalo reservoir is well established in South Africa, the FMDV situation is fundamentally different in that the World Organization of Animal Health (OIE) classifies southern Africa as a region of sporadic FMD outbreaks as opposed to the hyper-endemic status of FMDV in East Africa. Furthermore, buffalo in South Africa are largely separated from cattle by fencing (Dion et al., 2011, Miguel et al., 2013), which is in stark contrast to the extensive wildlife-livestock sympatry observed in semi-arid pastoralist rangelands in East Africa. Only SAT1 and 2 serotypes have been isolated from buffalo in Kenya, and the single sequences available each for buffalo-origin SAT1 and SAT2 viruses are genetically distinct from those in livestock, leading to the theory of independent virus populations in East African buffalo and cattle (Wekesa et al., 2015b). However, more genetic data on sequences found in buffalo are needed to elucidate the molecular epidemiology of FMDV in East African wildlife-livestock interfaces and the phylogenetic relationship between buffalo-derived viruses and vaccine strains.

Here, we assessed FMDV infection patterns in sympatric buffalo and cattle in central Kenya to evaluate the occurrence and dynamics of these viruses. We first investigated serological evidence of exposure to FMDV in buffalo and cattle. Second, we investigated the phylogenetic relationships of FMDV between cattle and buffalo with an aim to elucidate the extent to which transmission of FMDV occurs at wildlife-livestock interfaces in Kenya. These results further our understanding of the dynamics of FMDV in ecosystems characterized by mixed-grazing livestock and wildlife.

## Materials and Methods

### Description of the Study site

This study was conducted in Ol Pejeta Conservancy (OPC) in the Laikipia-Samburu ecosystem in central Kenya (Figure 1). The conservancy is situated along the equator at longitude 36.7°E, with a vegetation primarily comprised of grasslands interspersed with acacia, *Euclea* and riverine woodlands. The region has a bimodal rainfall pattern, with the long rainy season occurring in March-April, and the short rainy season in October-November. The average rainfall in this area is ~900mm per year (Birkett, 2002), though the running average across recent years (2004-2014) has been lower at ~600mm (OPC Ecological Monitoring Unit, unpublished). OPC is a mixed operation enterprise focusing jointly on cattle ranching and wildlife conservation. The conservancy has ~8,500 cattle, which are grazed during the day and stockaded in corrals during the night. The corrals are moved often based on management’s appraisal of pasture availability and ecological preservation of the grazing areas (VanderWaal et al., 2017). OPC has ~1,200 buffalo in herds of between 30-200 individuals. The Conservancy is fenced from the surrounding human settlements, but the fence has several openings to allow animal movement and maintain contiguity with the larger Laikipia-Samburu-Isiolo ecosystem. The buffalo in OPC are non-migratory. There is no separation of cattle, buffalo, and other wildlife, and these species are in continual spatial and temporal contact during grazing and at watering points. The Ewaso Ngiro River runs through the center of OPC, forming a natural barrier to animal movement. Cattle in OPC are first vaccinated when they are <12 months in age. OPC experienced a clinical outbreak of FMD in 2014.

**Figure 1:**
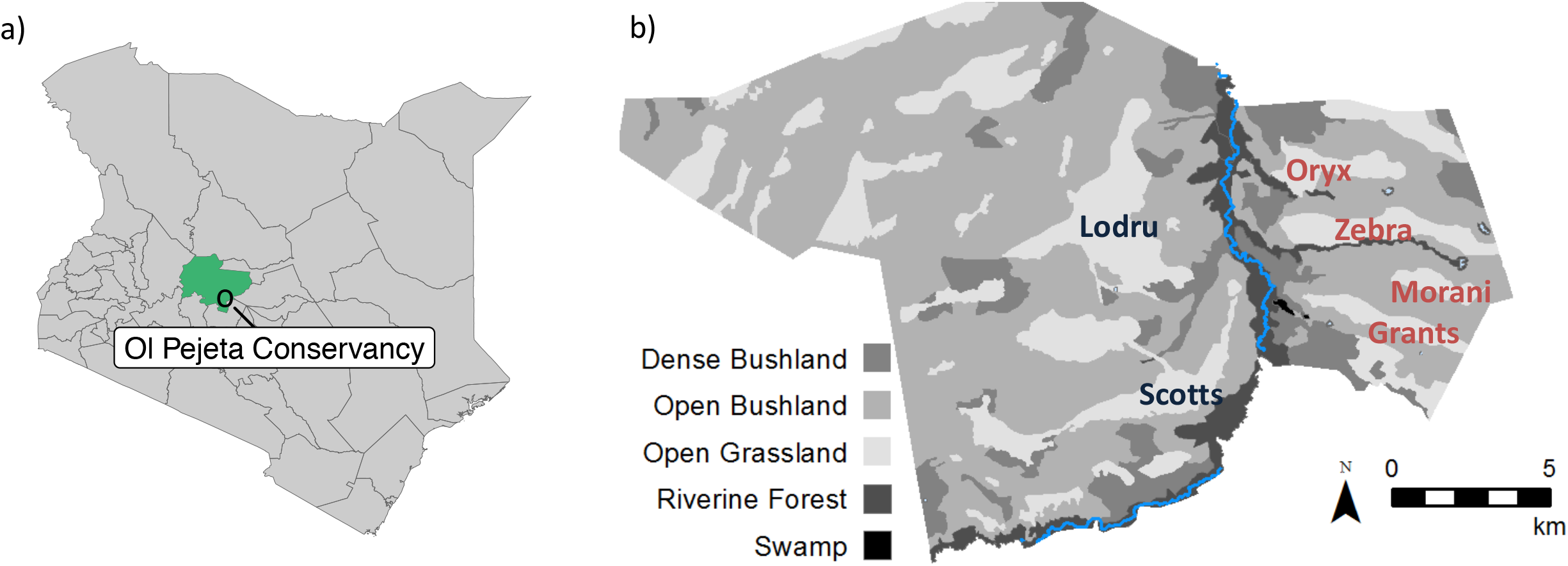
**a)** Map of Kenya showing location of the study site (O) within Laikipia County (green). **b)** Map of Ol Pejeta and locations of sampled buffalo herds.

### Biological sample collection

Buffalo were chemically restrained by Kenya Wildlife Service veterinarians using a combination of etorphine and azaperone delivered from a vehicle using a dart gun. All animals were sampled within five minutes of restraint to minimize negative welfare outcomes. Buffalo anesthesia was reversed using an antagonist (diprenorphine). Cattle were physically restrained. 5ml of blood was collected from the jugular vein into serum separator tubes, centrifuged, and harvested sera placed in cryovials for immediate preservation in liquid nitrogen. At the same time (during restraint), a probang cup was inserted into the oropharynx to draw out oropharyngeal fluids (OPF) from buffalo and cattle (Sutmoller and Gaggero, 1965). The fluid was aliquoted into cryovials containing either RNALater or Dulbecco’s Modified Eagle’s Medium (DMEM). The labeled samples were immediately preserved in liquid nitrogen and cold-chain was maintained during shipment until analysis.

Blood and OPF samples, together with associated metadata (sex, age, herd identification, sampling location and health status), were collected from 92 buffalo and 98 cattle (Table 1). Buffalo were sampled from nine distinct herds, and herd ID was based only on aggregations that existed at the time of sampling. While buffalo herds are known to exhibit fission-fusion dynamics (Cross et al., 2005b), longer-term behavioral data was not available to ascertain the permanence of observed herds. Because buffalo are typically infected at a young age and the quantity of virus that can be recovered declines over time (Leforban, 2010, BronsVoort et al., 2008) younger animals (<3 years) were preferentially selected to maximize the likelihood of recovering high-quality samples for genetic analysis. Cattle samples were collected from four OPC herds, and two sites located in human settlements adjacent to OPC. Because OPC experienced an FMDV outbreak in 2014, OPC herds were selected to capture younger animals that would have been alive at the time of the outbreak. In addition, this study included 21 cattle FMDV outbreak samples (vesicle epithelium or tissue homogenates, Table 2) collected by the FMD-L between 2014 and 2016 as part of their mandate. All procedures and protocols were ethically reviewed and approved. This study was approved by the Institutional Animal Care and Use Committee of the University of Minnesota (Protocol ID: 1502-32343A), and the Kenya Wildlife Service (KWS/BRM/5001). Outbreak samples and aliquots of all OPF samples were exported to Plum Island Animal Disease Center, USA under the approval of Directorate of Veterinary Services (RES/POL/VOL.XXVII/334) and the non-CITES permit (0004401).

**Table 1:**
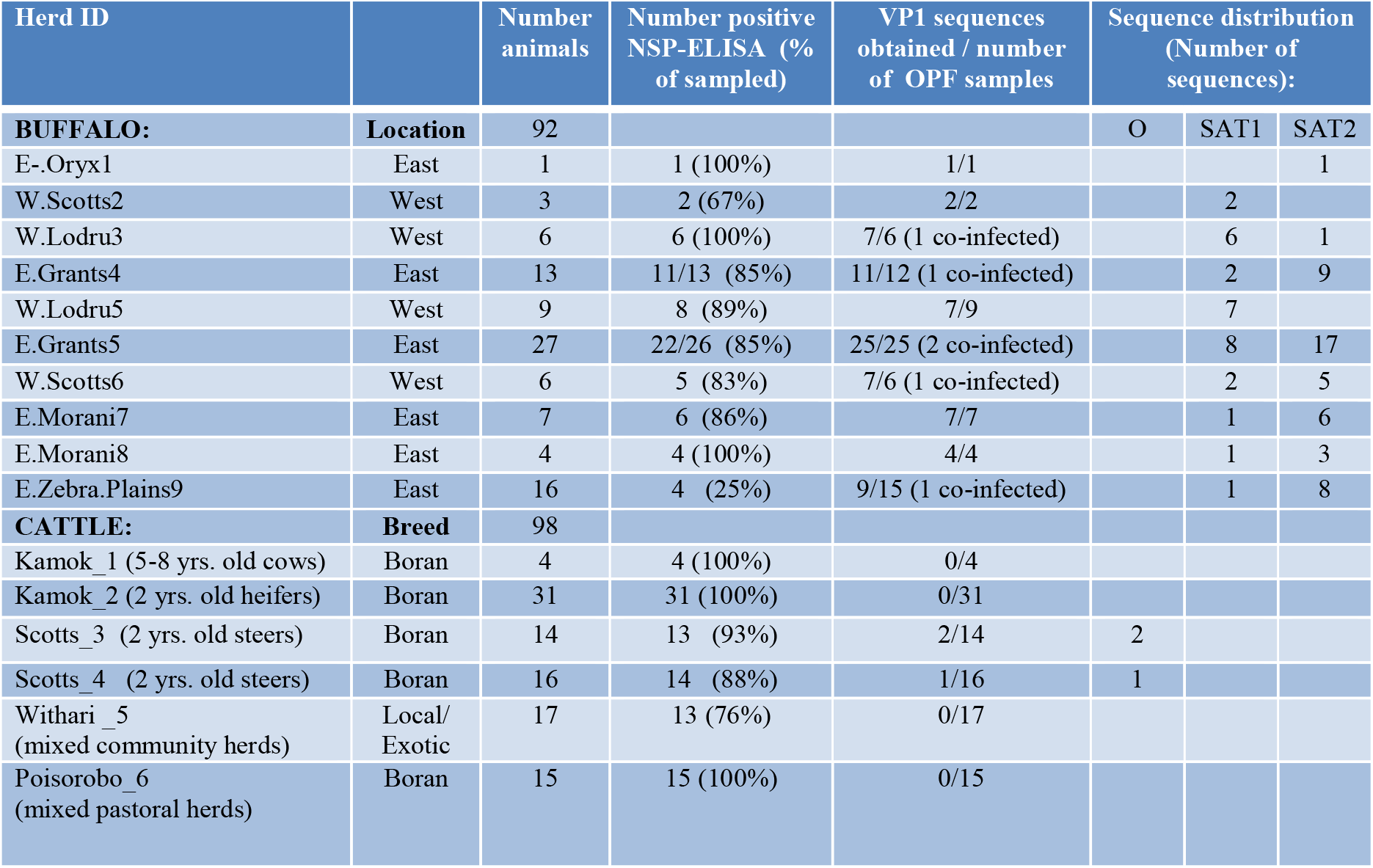
Summary of results, including number of animals sampled in each herd, number positive to FMDV by NSP-ELISA, number of VP1 sequences obtained, and the proportion of samples where each serotype was identified.

| Herd ID |  | Number animals | Number positive NSP-ELISA (% of sampled) | VP1 sequences obtained / number of OPF samples | Sequence distribution (Number of sequences): |  |  |
| --- | --- | --- | --- | --- | --- | --- | --- |
| **BUFFALO:** | **Location** | 92 |  |  | O | SAT1 | SAT2 |
| E.-Oryx1 | East | 1 | 1 (100%) | 1/1 |  |  | 1 |
| W.Scotts2 | West | 3 | 2 (67%) | 2/2 |  | 2 |  |
| W.Lodru3 | West | 6 | 6 (100%) | 7/6 (1 co-infected) |  | 6 | 1 |
| E.Grants4 | East | 13 | 11/13 (85%) | 11/12 (1 co-infected) |  | 2 | 9 |
| W.Lodru5 | West | 9 | 8 (89%) | 7/9 |  | 7 |  |
| E.Grants5 | East | 27 | 22/26 (85%) | 25/25 (2 co-infected) |  | 8 | 17 |
| W.Scotts6 | West | 6 | 5 (83%) | 7/6 (1 co-infected) |  | 2 | 5 |
| E.Morani7 | East | 7 | 6 (86%) | 7/7 |  | 1 | 6 |
| E.Morani8 | East | 4 | 4 (100%) | 4/4 |  | 1 | 3 |
| E.Zebra.Plains9 | East | 16 | 4 (25%) | 9/15 (1 co-infected) |  | 1 | 8 |

|  |  |  |  |  |  |
| --- | --- | --- | --- | --- | --- |
| **CATTLE:** | **Breed** | 98 |  |  |  |
| Kamok_1 (5-8 yrs. old cows) | Boran | 4 | 4 (100%) | 0/4 |  |
| Kamok_2 (2 yrs. old heifers) | Boran | 31 | 31 (100%) | 0/31 |  |
| Scotts_3 (2 yrs. old steers) | Boran | 14 | 13 (93%) | 2/14 | 2 |
| Scotts_4 (2 yrs. old steers) | Boran | 16 | 14 (88%) | 1/16 | 1 |
| Withari_5<br>(mixed community herds) | Local/<br>Exotic | 17 | 13 (76%) | 0/17 |  |
| Poisorobo_6<br>(mixed pastoral herds) | Boran | 15 | 15 (100%) | 0/15 |  |

**Table 2:** Summary of outbreak samples for which sequences were obtained; outbreak occurring in OPC is bolded.

| ID | Serotype | Topotype | Location | Sub-county | County | Date |
| --- | --- | --- | --- | --- | --- | --- |
| K14/14 | SAT1 | I(NWZ) | Kirimene | North Imenti | Meru | 10-Jan-14 |
| K19/14 | SAT1 | I(NWZ) | Mochongoi | Baringo | Baringo | 16-Jan-14 |
| K29/14 | SAT1 | I(NWZ) | Kabiru-ini | Nyeri Central | Nyeri | 29-Jan-14 |
| K60/14 | O | EA_2 | Subukia East | Subukia | Nakuru | 5-Mar-14 |
| K103/14 | A | Africa_GI | Gotu | Isiolo | Isiolo | 2-Jul-14 |
| K137/14 | SAT2 | IV | Rumuruti | Rumuruti | Laikipia | 11-Sep-14 |
| **K145/14** | **A** | **Africa_GI** | **Ol Pejeta** | **Laikipia East** | **Laikipia** | **20-Sep-14** |
| K39/15 | A, SAT2 | Africa_G1 / IV | Losogwa | Nyahururu | Laikipia | 12-Mar-15 |
| K91/15 | A |  | Nyambogichi | Laikipia East | Laikipia | 26-Sep-15 |
| K103/15 | A |  | Tulasha | Nyandarua | Nyandarua | 31-Oct-15 |
| K19/16 | O | EA_2 | Rurii | Ol Kalau | Nyandarua | 20-Apr-16 |
| K74/16 | A | Africa_GI | Subukia East | Subukia | Nakuru | 13-Oct-16 |
| K75/16 | SAT1 | I(NWZ) | Munyeke | Nyandarua | Nyandarua | 27-Oct-16 |

### Serological analysis

OPC cattle were usually vaccinated against various diseases during their first year of age, including FMDV. Thus, to discriminate against vaccine versus natural FMDV infection, we analyzed serum samples for response against FMDV non-structural proteins (NSP-ELISA, Priocheck^®^, Prionics AG, Netherlands) following the manufacturer protocol. Briefly, both buffalo and cattle serum samples were heat-inactivated then added into the 3ABC specific coated plates and incubated at 22o Celsius overnight. The plates were then washed using 100μL of the conjugate and further incubated at the same temperature for an hour. The plates were then subjected to another round of washing and 100μL of Chromogen substrate added to each well. This procedure was followed by 30 minutes incubation. After adding the stop solution, the optical density was measured at 450nm where a sample was considered positive for FMDV if the percentage inhibition was greater than fifty percent.

### Molecular analysis of viral RNA

#### FMDV RNA detection and sequencing

All OPF and vesicle epithelium samples were screened for the presence of FMDV RNA using real-time reverse transcriptase polymerase chain reaction (rRT-PCR) as previously described (Arzt et al., 2010, Pacheco et al., 2010). Briefly, RNA was extracted using either the RNeasy mini kit (Qiagen, Hilden, Germany) or the MagMAX^™^-96 viral RNA isolation kit (Ambion, Austin, TX, USA) following manufacturers’ instructions. RNA was analyzed by rRT-PCR following a previously described protocol (Pacheco et al., 2010, Callahan et al., 2002). Samples were considered positive when *C*t values were <40. The viral RNA from positive samples was further analyzed by RT-PCR and sequenced using the Sanger method as previously described (Ludi et al., 2016) or by next generation sequencing (NGS). Briefly, for NGS-derived sequences, RT-PCR amplicons were produced using the universal FMDV primers described by Xu *et al*. (2013), analyzed and purified as previously described (Ludi et al., 2016). The purified amplicons were processed with the Nextera XT DNA library kit (Illumina, Catalog #FC-131-1096) and sequenced on an Illumina NextSeq 500.

Sequence reads from each sample were mapped to a set of reference published sequences of all serotypes. We identified the sequence to which the majority of the reads mapped and *de novo* assembled the mapped read to obtain the consensus sequence. If a high number of reads were assembled to two different reference serotypes, the reads that mapped to each serotype were obtained, mapped to the corresponding reference and the consensus obtained. Subsequently, the consensus sequence obtained by the previous step was queried in BLAST (https://blast.ncbi.nlm.nih.gov/) to identify if a more suitable reference was available to obtain the final consensus sequence. If a more suitable reference was identified, the reads were mapped to it, and the final consensus sequence was obtained. In general, over 99% of the NGS reads from each individual sample mapped to a single reference sequence. However, two distinct serotypes (SAT1 and SAT2) were concurrently found in six individual buffalo OPF samples. For these six OPF samples, ~30-70% of the reads mapped to a SAT1 reference sequence while the remaining NGS reads mapped to a SAT2 reference sequence. Final sequences used in this study covered 100% of the VP1 coding segment and had a minimum depth of coverage of >100. All assembly steps were performed in CLC Workbench 10.

#### Statistical and phylogenetic analysis

At sampling, metadata were collected from each animal, including sex, age, herd identification, and sampling location. Descriptive statistics were tabulated, including summary of number of animals sampled in each herd, number FMDV-positive by NSP-ELISA, and number of VP1 sequences obtained. Generalized linear models (GLMs), performed in R v3.4.2 (www.R-project.org) were used to evaluate the effect of sex, herd size, and sampling location on the likelihood of being positive for FMDV.

Phylogenetic analysis was conducted on each FMDV serotype recovered in this study. Additional viral VP1 sequences were sourced from GenBank using BLAST, implemented in Geneious v11.1.4 (https://www.geneious.com) and aligned using MUSCLE. Phylogenetic trees were constructed for each serotype using maximum likelihood with the GTR-G nucleotide substitution model and 1000 bootstraps, implemented in RAxML v8 (Stamatakis, 2014). Resulting trees were edited using Figtree 1.4.3 (http://tree.bio.ed.ac.uk/software/figtree/) and Tree Graph 2 (Stover and Muller, 2010).

In East Africa, the Kenya Veterinary Vaccine Production Institute produces a polyvalent FMD vaccine which includes serotypes O, A, SAT1 and SAT2. Percent identity was calculated at the nucleotide and amino acid levels to compare new sequences and vaccine strains using Geneious (Serotype A (strain/NCMI accession number): K35/80-KJ440846 and K5/80-KJ440848; O: K77/78-HM756588; SAT1: T155/71-HQ267519; SAT2: K52/84-HM623685).

To assess whether the FMDV phylogenetic trees exhibited any clustering according to the buffalo population’s social (according to herd) or spatial structure (according to the herd’s location in OPC: east or west of the river), we applied analytical methods developed in community ecology as a means to use phylogenetic data to infer local ecological (or epidemiological) processes (Fountain-Jones et al., 2018, Tucker et al., 2017). Specifically, we used a metric of phylogenetic divergence (i.e., beta-diversity), COMDIST, that quantifies the extent of dissimilarity amongst the FMDV sequences from each buffalo community (defined as herd or side of river)(Webb et al., 2008). COMDIST is defined as the mean pairwise phylogenetic distance among sequences found in two different communities and was calculated for each pair of communities using the R package *pez* (Pearse et al., 2015). Herds with only one or two sequences were excluded due to low sample size. Therefore, only three herds were included for SAT1 (Herd IDs: E.Grant5, W.Lodru3, and W.Lodru5), and six herds for SAT2 (Herd ID: E.Grants4, E.Grants5, E.Morani7, E.Morani8, E.Zebra.plains9, W.Scotts6, Table 1). In addition, outlying sequences were excluded otherwise their pairwise phylogenetic distances from other sequences disproportionately influenced COMDIST calculations (PRB84 for SAT1; PRB13, PRB50, and PRB75 for SAT2). We performed Monte Carlo permutations to determine if the observed divergence values were statistically non-random or whether they could have emerged by chance. We performed constrained randomizations where buffalo sample IDs were reshuffled, effectively creating a null distribution of the expected divergence values if the virus sequences found in each buffalo were independent of community identity. For each pair of communities, a p-value was calculated as the number of permutations that produce divergence values more extreme than the observed value. Significance tests were two-sided and Bonferroni corrections were made to account for multiple pairwise comparisons. Monte Carlo p-values can be interpreted as the probability that the observed divergence among communities would occur if community structure was independent of phylogenetic structure. In other words, low p-values indicate that virus transmission between communities is infrequent.

## Results

### FMDV serology

Blood, sera, and OPF samples were collected from 98 asymptomatic cattle and 92 buffalo. In addition, 21 samples were available from previous outbreaks in cattle. 93% of the cattle sampled (95% CI: 88 – 98) had anti-FMDV-NSP antibodies, while 77% (95% CI: 66 – 85) of sampled buffalo were positive for anti-FMDV-NSP. Neither sex, herd size, nor sampling location were significantly associated with a herd of buffalo or cattle being positive for FMDV.

### VP1 Sequences and Phylogenetic analysis

We recovered 80 VP1 sequences from the buffalo OPF samples, all of which were serotype SAT1 (30 sequences) and SAT2 (50 sequences). Though several herds in OPC appeared to exclusively have serotype SAT1 (Table 1), SAT2 was more common in most herds. In addition, six individual buffalo had dual infections with both SAT1 and SAT2 serotypes. Further, we recovered 14 VP1 sequences from cattle sampled during outbreaks and three VP1 sequences from cattle OPF samples collected at OPC. Outbreak samples were from serotypes SAT1, SAT2, O and A (Table 2). All the recovered sequences from cattle OPF (this study) were serotype O.

Depending on serotype, percent identities between new VP1 sequences and vaccine strains used in Kenya ranged from a mean of 82 to 89% at the nucleotide level and 87% to 94% at the amino acid level (Table 3). The lowest mean percent identity found for serotype A and the highest for SAT1. Interestingly, buffalo sequences were more similar to the SAT2 vaccine (87% nucleotide identity, 92% amino acid identity, Table 3) than were cattle (78% nucleotide identity and 87% amino acid identity). Such differences amongst cattle and buffalo relative to the vaccine were not observed for SAT1 (Table 3).

**Table 3:** Percent nucleotide and amino acid similarity of new sequences compared to vaccine strains (Mean and standard deviation – SD).

|  | Nucleotide | Amino Acid |
| --- | --- | --- |
|  | Mean (SD) | Mean (SD) |
| A | 82.2 (0.8) | 87.4 (3.5) |
| O | 84.3 (0.4) | 92.4 (0.5) |
| SAT1<br>(all) | 89.8 (0.9) | 94.2 (1.3) |
| SAT1<br>(buffalo) | 89.7 (0.8) | 94.1 (1.4) |
| SAT1<br>(cattle) | 90.9 (0.8) | 95.0 (0.7) |
| SAT2<br>(all) | 86.3 (3.4) | 92.0 (2.4) |
| SAT2<br>(buffalo) | 87.0 (0.9) | 92.5 (1.1) |
| SAT2<br>(cattle) | 78.6 (7.6) | 87.6 (0.7) |

Five serotype A sequences were recovered from cattle outbreak samples, one of which as the virus associated with the FMD outbreak in OPC in 2014 (K145). The OPC outbreak was genetically closely related to 2014 and 2015 outbreaks occurring in neighboring counties (Isiolo and Nyahururu). Two other outbreak samples that were analyzed as part of this study were more closely related to those from Tanzania and Uganda than to those from Kenya (Figure 2). We recovered five serotype O sequences, all from cattle samples. One recovered sequence was from a cattle outbreak in Nyandarua, a neighboring county to Laikipia County. This sequence was closely related to those from Tanzania and other parts of Kenya from outbreaks in 2010. Three additional sequences were analyzed as part of this study from other cattle outbreaks in Kenya in 2010. These sequences were closely related to one of our study’s samples sequences recovered from a 2014 cattle outbreak in Nakuru County, Kenya. These sequences were closely related to samples collected during outbreaks in the former Rift Valley and Central provinces in 2011 in Kenya (Figure 3).

**Figure 2:**
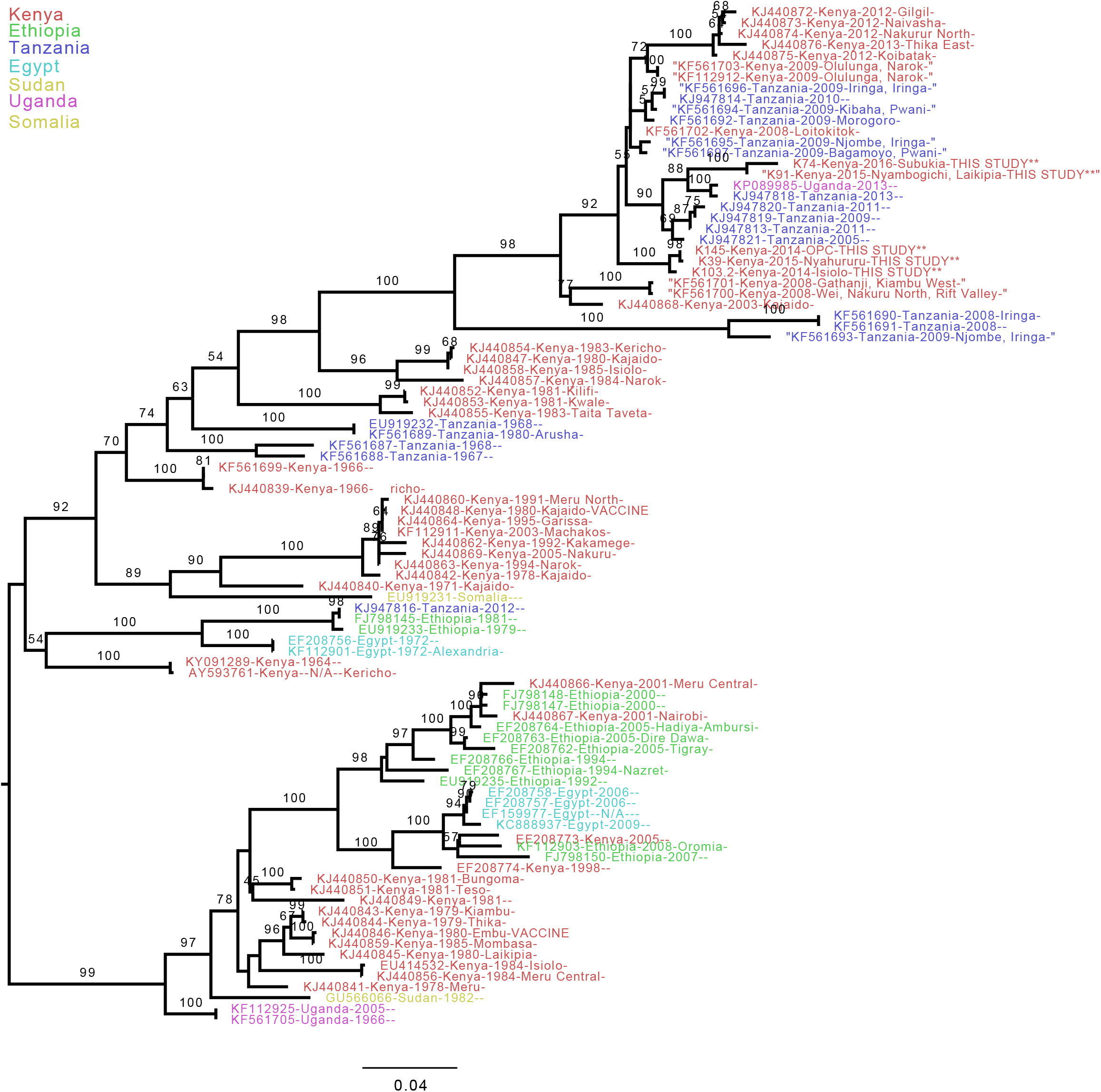
Phylogenetic relationship of FMDV serotype A reconstructed using maximum likelihood methods with 1000 bootstraps. The vaccine strains and sequences recovered from this study are noted in capital letters. The five sequences from our study, all from cattle outbreaks are shown (location of sampling noted in tip labels).

**Figure 3:**
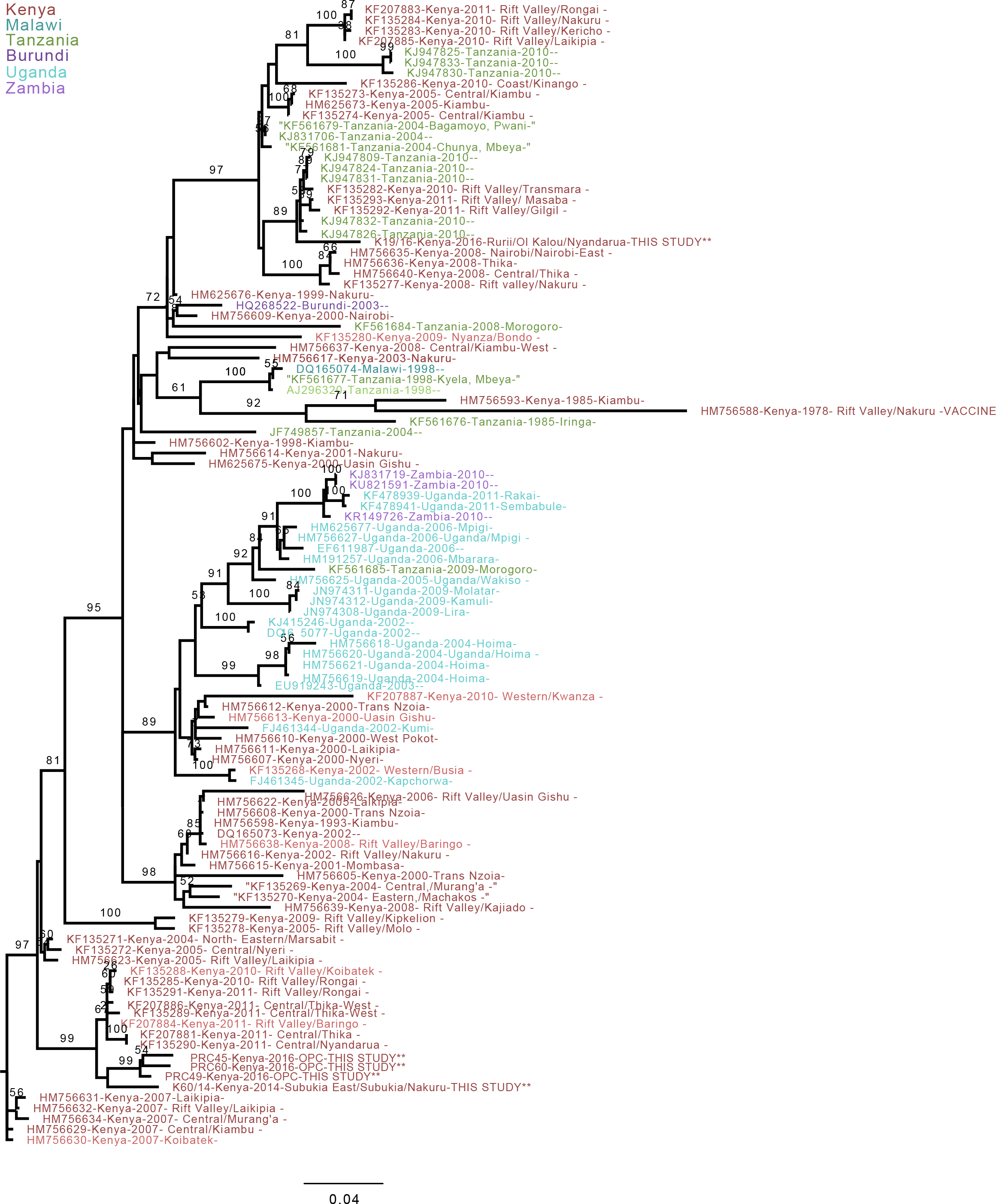
Phylogenetic relationship of FMDV serotype O reconstructed using maximum likelihood methods with 1000 bootstraps. The colors represent country from where the sequences were recovered. The vaccine strains and sequences recovered from this study are noted in capital letters. The five sequences from our study, all from cattle outbreaks are shown (location of sampling noted in tip labels).

We recovered SAT1 sequences from 30 buffalo and four from cattle outbreak samples (Figure 4). Sequences recovered from cattle outbreaks in Samburu and Nyeri counties in 2014 were found to be closely related to buffalo sequences PRB25, PRB26 and PRB27, whereas a sequence from an outbreak in 2010 from Laikipia County was closely related to sequences PRB3 and PRB4 from our study. The only buffalo SAT1 sequence from a previous study, isolated from buffalo in Tsavo East National Park located ~650 km away, was closely related to our buffalo FMDV sequences.

**Figure 4:**
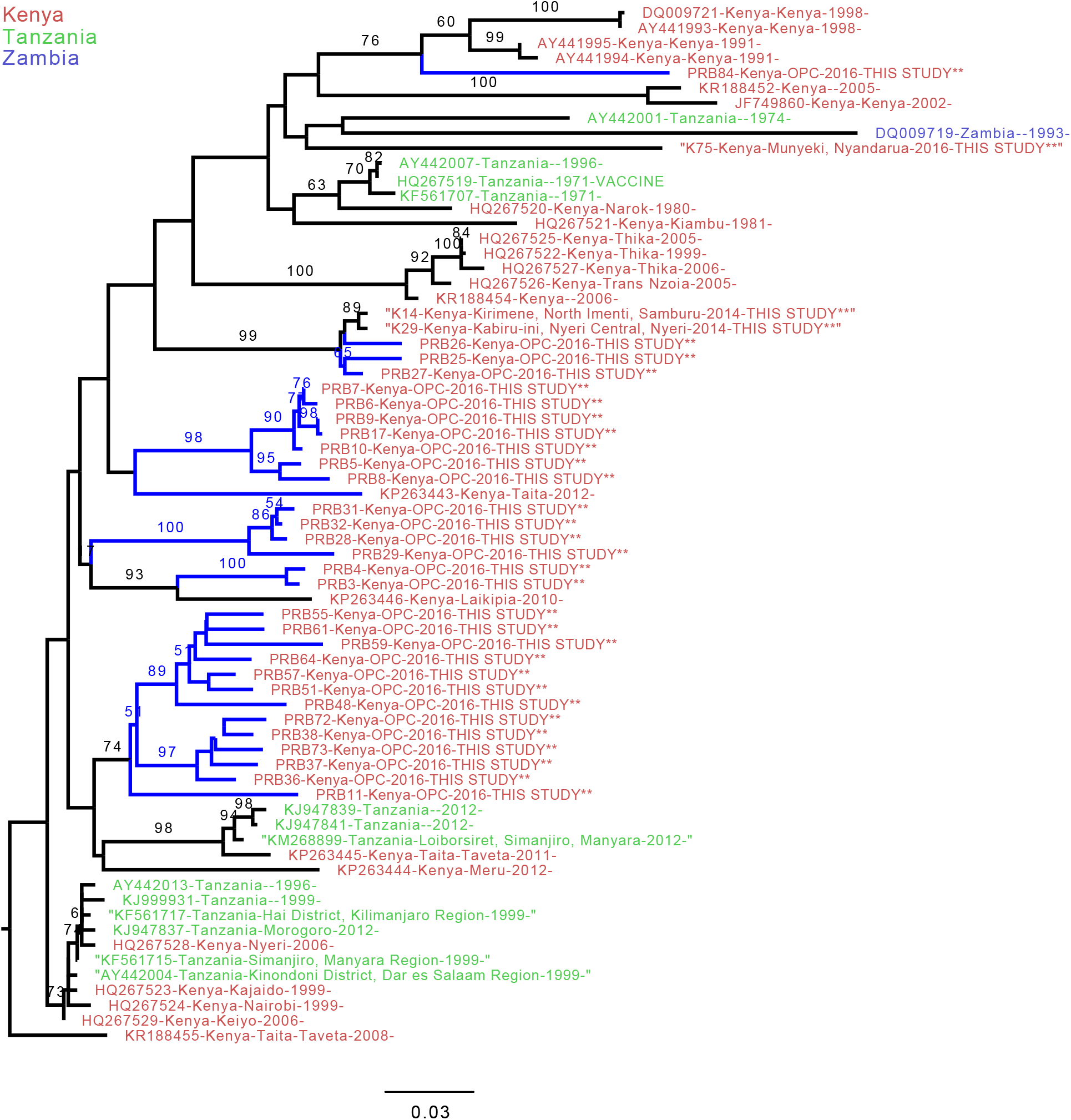
Phylogenetic relationship of FMDV serotype SAT1 reconstructed using maximum likelihood methods with 1000 bootstraps. The colors represent the country, while vaccine strains and sequences from this study are noted in capital letters. Thirty-two sequences from our study are shown with three sequences coming from cattle outbreaks. Branches marked in blue are from buffalo OPF samples (all but one of which come from this study).

We recovered SAT2 sequences from 50 buffalo and two cattle outbreaks (Figure 5). The single buffalo SAT2 sequence previously available was from Narok County, and was not closely related to the buffalo sequences from this study, whereas sequences from cattle outbreaks in Rumuruti and Nyahururu (towns adjacent to our study area) in 2014 and 2015 appear somewhat closely related to the buffalo sequences from our study. Interestingly, it appears that there is more variability in SAT2 than in SAT1 sequences recovered from buffalo populations in this study. Additionally, SAT2 sequences from buffalo and cattle appear less inter-mixed than SAT1.

**Figure 5:**
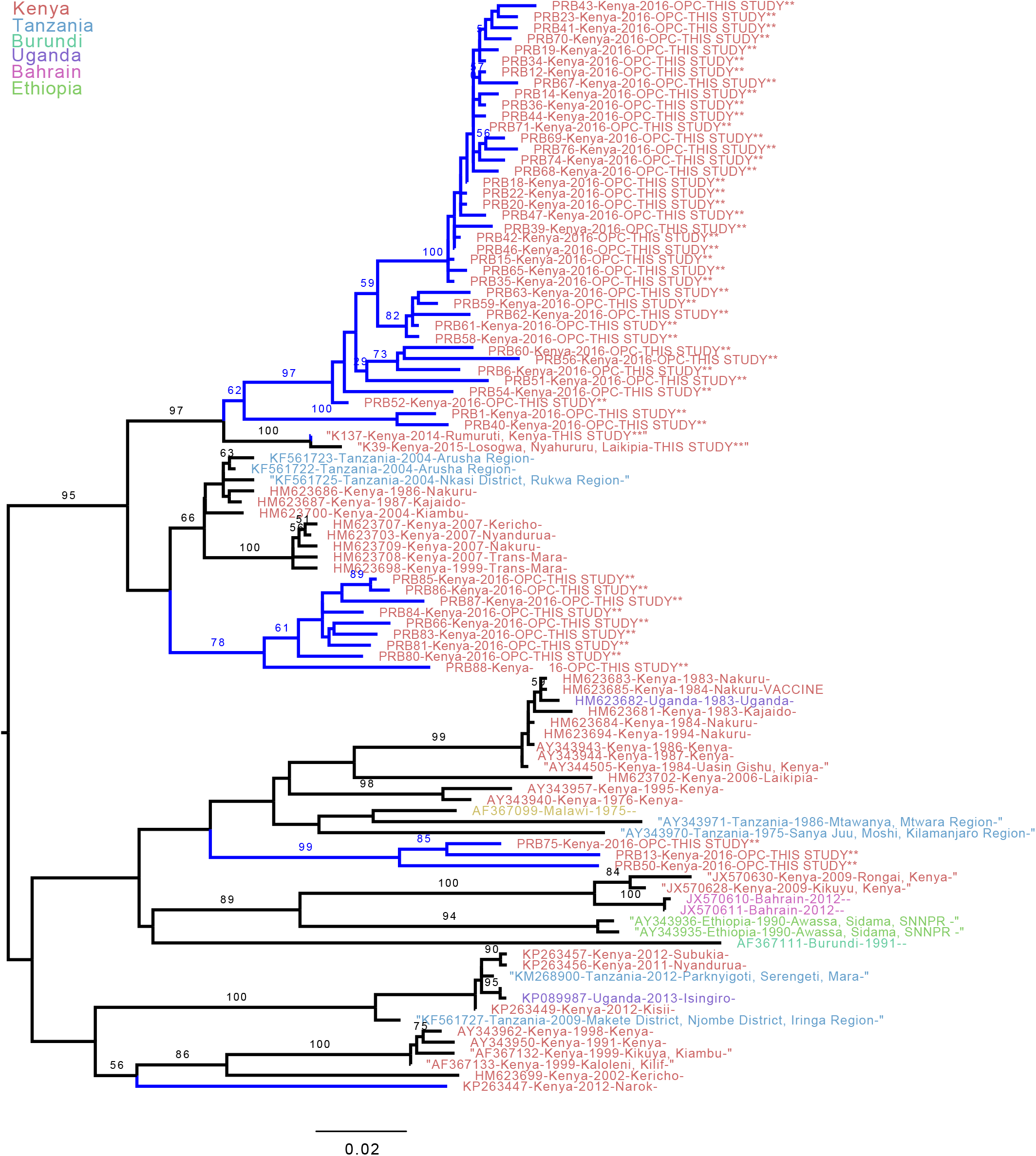
Phylogenetic relationship of FMDV serotype SAT2 reconstructed using maximum likelihood methods with 1000 bootstraps. The colors represent country, while vaccine strains and sequences from this study are noted in capital letters. Fifty-three sequences from our study are shown. Two sequences come from cattle outbreaks. Branches marked in blue are from buffalo OPF samples (all but one of which come from our study).

Ol Pejeta Conservancy is subdivided by the Ewaso Ngiro River (Figure 1b), which forms a natural semi-permeable barrier to animal movement. Based on the SAT1 phylogenetic tree, sequences from the buffalo sampled from herds on the east versus west side of the river cluster separately, with significant phylogenetic divergence between those two communities of buffalo (p < 0.001, Figure 6a). This pattern is not as apparent for SAT2 (p < 0.26, Figure 6b), perhaps because SAT2 is relatively rare on the west side. For both serotypes, phylogenetic divergence was also present at the herd level. All three herds included in the SAT1 analysis had significant phylogenetic divergence from one another (p < 0.001 for all pairwise comparisons of herds, Figure 6a, 7a). SAT2 phylogenetic divergence was largely related to the distinct genetic variant prevalent in the Zebra Plains herd (p < 0.005 for all pairwise comparisons with E-Zebra.Plains9, Figure 6b, 7). This variant accounted for 8 of 9 sequences recovered from this herd, but was rare or absent in all other herds. For SAT2, several herds had VP1 sequences that were significantly less divergent than expected by chance. Specifically, E-Grants4 was clustered with E-Grants5, E-Morani7, and E-Morani8 (p < 0.001 for pairwise comparisons with E.Grants4, Figure 6b, 7).

**Figure 6.**
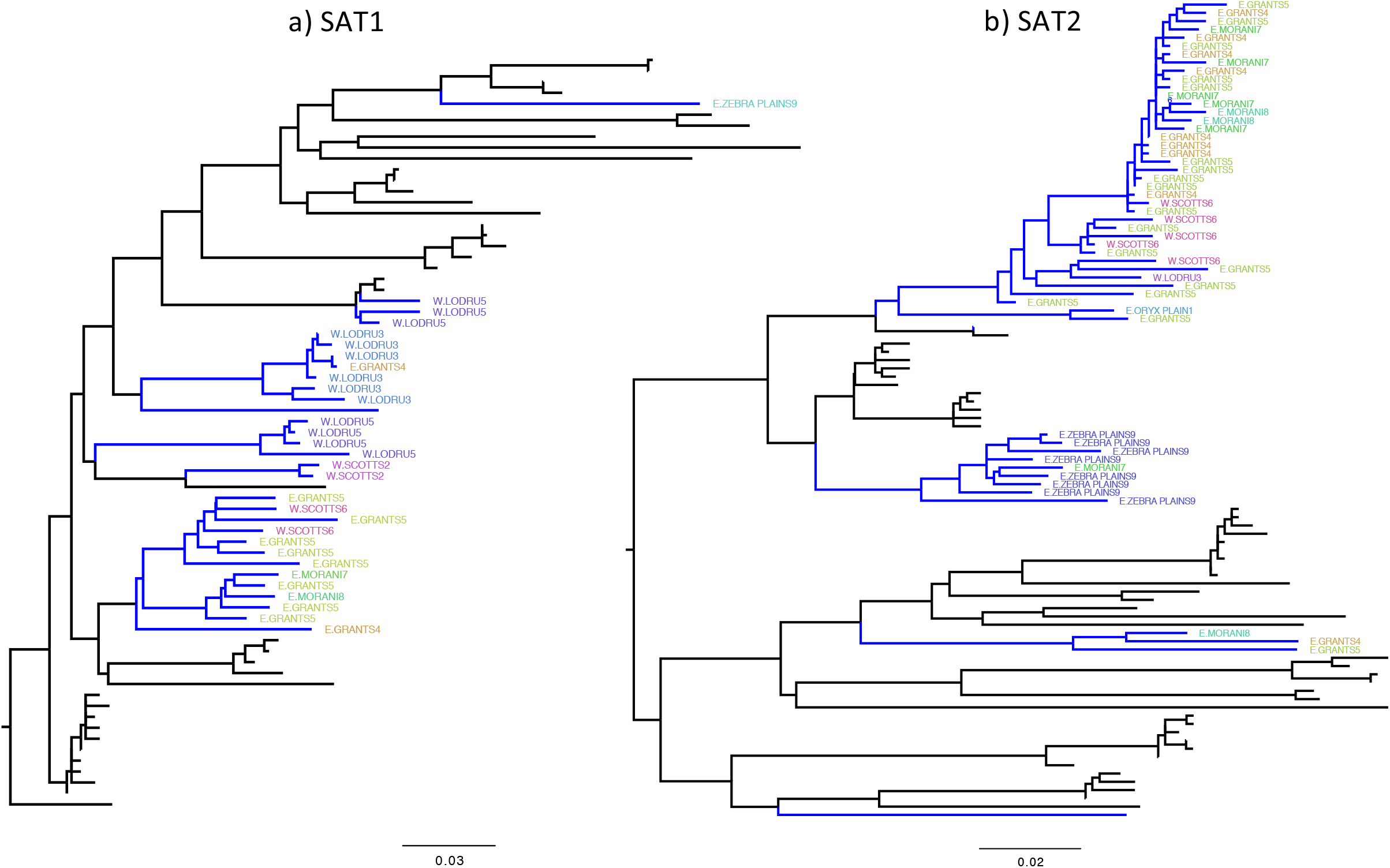
Phylogenetic trees for a) SAT1 and b) SAT2, where label colors represent herd. Herd ID prefixes E versus W indicates whether the herd was sampled on the east or west side of Ewaso Ngiro River that sub-divides the study area. The branches marked in blue are from buffalo OPF samples.

**Figure 7.**
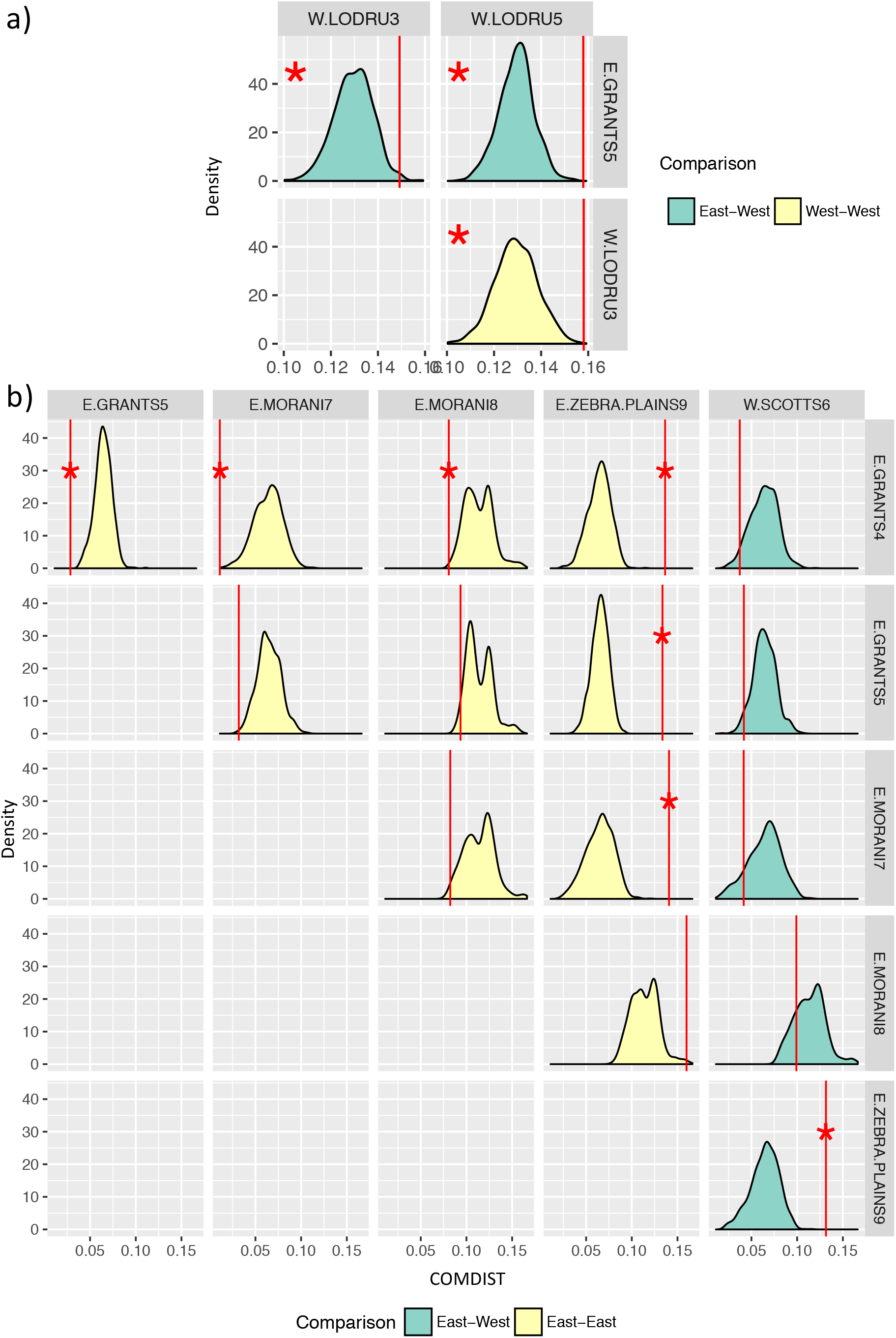
Phylogenetic divergence at the herd-level for a) SAT1 and b) SAT2 FMDV isolated from buffalo in OPC. Vertical red lines represent the observed value of divergence (COMDIST) between each pair of herds. Shaded regions represent the null density distribution of divergence values (COMDIST) if herd membership is randomized. The color of the shading indicates whether each pair of herds was on the same or different sides of the river. Stars indicate that the observed value is significantly different from null expectations.

## d) Discussion

In this study, we used molecular and serological analyses to investigate FMDV infection patterns at the wildlife-livestock interface in Kenya. We found that four serotypes (A, O, SAT1, and SAT2) were co-circulating within a single ecosystem in central Kenya. While we expected that a highly contagious disease such as FMDV would be readily transmitted among sympatric host species, we did not find evidence for buffalo-cattle transmission within the OPC ecosystem. However, serotype SAT1 and SAT2 sequences associated with FMDV outbreaks in cattle elsewhere in the region were phylogenetically related to buffalo sequences, suggesting that cross-species transmission does occur and that FMDV found in buffalo and cattle do not represent independent viral populations. We also showed that transmission of FMDV in buffalo populations was influenced by fine-scale social and spatial structure. Although we cannot confirm the directionality of cross-species transmission from this analysis, these results indicate that buffalo may play an important role in the epidemiology of FMDV in Kenya.

For all FMDV serotypes, the vaccine strain was not closely related to the sequences isolated from Kenya in this study (Table 3), with percent nucleotide and amino acid identity never exceeding 89% and 94%, respectively. However, percent identity is not a perfect measure of vaccine performance and antigenic matching was beyond the scope of this study. Interestingly, the buffalo SAT2 sequences were more similar to the vaccine strain than SAT2 viruses from cattle. This perhaps suggests that some cattle SAT2 viruses either have been introduced from outside of Kenya or have undergone more rapid evolution as might be expected in outbreak situations (Pedersen et al., 2015).

The 2014 outbreak of FMDV in OPC was caused by serotype A. The VP1 sequence associated with this outbreak was closely related to viruses from outbreaks in 2014 and 2015 in Nyahururu and Isiolo (Figure 6), both of which are located relatively nearby. OPC sometimes purchases animals to fatten for slaughter from communities in the Isiolo area, which suggests a likely epidemiological linkage via animal movements between OPC and the Isiolo area. No serotype A viruses were recovered from OPC cattle OPF samples collected in 2016 despite the fact that these animals were alive during the previous outbreak. Such findings are not surprising given the fact that the carrier stage, during which cattle may show persistent infection in the nasopharynx, usually lasts only one year (Hayer et al., 2018, Bronsvoort et al., 2016) and up to 28 months under some conditions (Bertram et al., 2018). The only serotype to be recovered from OPC cattle OPF samples was serotype O, which was closely related to an outbreak of serotype O in Subukia, Nakuru, and another location from which OPC occasionally purchases animals. The presence of serotype O in OPC OPF samples was not associated with a reported clinical outbreak at OPC. Given lack of the individual histories of the three OPC cattle with serotype O, it is unclear how or when these animals became infected.

Two lines of evidence suggest that transmission between OPC buffalo and cattle is infrequent despite sympatry. First, no SAT1 and SAT2 virus was found in OPC cattle despite the high prevalence of SAT1 and SAT2 carriers in the buffalo population. Second, OPC cattle are relatively separated from neighboring community herds by a fence, and buffalo are rare or absent in the surrounding community. OPC cattle experience very few clinical outbreaks of FMDV compared to the surrounding communities, despite the fact that OPC cattle have a far higher amount of exposure to buffalo. This result suggests that the infection pressure due to circulation of FMDV in livestock populations is far higher than infection pressure from buffalo to cattle. Instead, the occurrence of FMDV in OPC cattle (serotypes O and A) was likely related to the introduction of animals from other locations or through human-associated fomites.

However, we did find substantial evidence that FMDV virus found in buffalo and cattle are not independent viral populations. Numerous cases of SAT1 and SAT2 virus associated with outbreaks in cattle were closely related to viruses present in the buffalo population. Prior to this study, only two sequences were available from buffalo in Kenya (Wekesa et al., 2015a). Interestingly, the only buffalo SAT1 sequence in Wekesa *et al*. (Wekesa et al., 2015a), which was collected in Tsavo East National Park, is closely related to viruses found in OPC buffalo despite that Tsavo is located over 650 km from OPC. The buffalo SAT2 sequence reported by Wekesa (Wekesa et al., 2015a) was not closely related to FMDV found in OPC buffalo. The eight SAT1 and five SAT2 sequences available from buffalo elsewhere in East Africa (Uganda) were different topotypes (Ayebazibwe et al., 2010b). For both SAT1 and SAT2, there were also several lineages found in OPC buffalo that were relatively rare (i.e., did not cluster with the other buffalo sequences) but were more closely related to historical outbreak sequences found in GenBank. From this analysis, we cannot determine whether buffalo were the source of these outbreaks in cattle, or if the buffalo acquire SAT1 and SAT2 virus already circulating in livestock. Future directions include a more quantitative evaluation of the frequency of cross-species transmission using Bayesian phylogenetic approaches. In addition, further sampling of wildlife populations across Kenya is needed to gain a more representative view of the diversity of FMDV found in buffalo in relation to outbreaks in livestock.

Interestingly, far greater numbers of SAT2 viruses (50 sequences) were recovered from the OPC buffalo population as compared with SAT1 (30 sequences), which is consistent with serological evidence regarding the prevalence of each serotype in buffalo in East Africa (BronsVoort et al., 2008). This is in contrast to southern Africa, where SAT1 is generally considered to be the dominant SAT serotype in buffalo based on both experimental studies (Maree et al., 2016) and field epidemiology (Bastos et al., 2001). Thus, it was surprising to see that SAT2 was 1.6x more common than SAT1 amongst the 80 buffalo FMDV sequences from this population. There are two potential explanations for this finding. First, it is possible that SAT2 behaves differently in East African than in southern Africa, where most of the previous research has been performed. Speculatively, this could reflect different patterns of host-pathogen co-evolution in southern versus eastern Africa after the FMDV SAT populations went through a population bottleneck during the great rinderpest epidemic in 1887-1897, which wiped out 95% of wild and domestic ruminants (Lasecka-Dykes et al., 2018, Casey et al., 2014). Alternatively, given that SAT2 appears to be better able to cause outbreaks cattle than SAT1 (Bastos et al., 2001, Bastos et al., 2003), the dominance of SAT2 in this East African buffalo population could reflect a higher SAT2 burden in cattle populations and subsequent spillover from cattle into buffalo.

Within the buffalo population studied here, FMDV exhibited significant phylogenetic divergence related to the population’s spatial (according to side of river) and social structure (according to herd). Buffalo herds tend to utilize defined home ranges for several years (Funston et al., 1994, Prins, 1996), which may contribute to the observed spatial and herd-level clustering of FMDV. For SAT1, significant divergence was observed that related to the presence of the river. One cluster was made up of four herds sampled to the east of the Ewaso Ngiro River (Grants and Morani herds), while the other cluster consisted primarily of animals sampled on the west side of the river (Lodru and Scotts herds). Animals do cross the river, and cattle are often moved from one side to the other. However, the river appears to delineate separate viral populations even though these buffalo are separated by only ~5 km, thus supporting lack of airborne or waterborne viral dispersal under these conditions. The apparent impact of the river on limiting transmission is consistent with what has been observed for *Escherichia coli* in wildlife in this population, where wildlife sampled on either side of the river were more likely to harbor genetically distinct *E. coli* (VanderWaal et al., 2014). This spatial pattern was not significant for SAT2 viruses.

Herd identity also influenced the phylogenies of SAT1 and SAT2 in buffalo, even for herds that were sampled from the same area (i.e., W.LODRU3 and W.LODRU5 for SAT1). This was particularly apparent for SAT2, where eight out of fifteen sampled animals in one herd (Zebra Plains) harbored a genetically distinct variant of FMDV. Buffalo sociality is characterized by fission-fusion dynamics, where herds come together and break apart over time (Cross et al., 2005a, Cross et al., 2005b). Thus, some of the herds sampled may not truly represent distinct social units but rather a snapshot of buffalo groupings at that time of sampling. For example, there was a lack of divergence of the SAT2 viruses between the herds sampled on Grants and Morani plains, suggesting that these herds may not be distinct social units. That being said, the divergence of the SAT1 viruses between the two Lodru herds and the distinctiveness of the FMDV found in the Zebra Plains herd suggests that some herds tend not to socialize with other nearby herds (Figure 1), and that social segregation may limit transmission.

Our results support the idea of maintenance of viruses within specific herds or areas, potentially through intergenerational transmission. FMDV can be maintained in an isolated buffalo herd for decades, suggesting that transmission occurs across generations within the herd (Cross et al., 2005a, Condy et al., 1985). The duration of persistent infection in carrier buffalo and the incidence of neoteric subclinical infection (Farooq et al., 2018) would be important for intergenerational transmission. These infection dynamics in subclinically infected animals would define the extent to which animals remain infectious long enough to allow for the birth of susceptible calves, thus maintaining a within-herd transmission cycle. Alternatively, the observed social and spatial clustering of viruses within the phylogenetic tree could also be explained by modularity of contact networks (Sah et al., 2017); the structure of contact networks between herds could lead to differential transmission opportunities and, consequently, to the herd-level differences observed in the phylogenetic trees. This seems less likely, given evidence from Kruger National Park, South Africa, in which buffalo-origin VP1 sequences clustered spatially but not temporally (Bastos et al., 2000b, Vosloo and Thomson, 2017), highlighting the potential importance of within-herd or within-area transmission cycles. Additional research focused on longitudinal and behavioral sampling would be necessary to fully elucidate transmission processes in East Africa.

In conclusion, evidence from the ecology of FMDV in the OPC ecosystem suggests that transmission of FMDV between cattle and buffalo is rare. Instead, the occurrence of FMDV in OPC cattle appears to be related to the introduction of cattle from other regions. However, viruses from outbreaks in cattle elsewhere in the country were caused by viruses closely related to SAT1 and SAT2 viruses found in buffalo. Thus, the circulation of FMDV in buffalo and the sharing of FMDV between wildlife and livestock must be addressed in order to decrease the occurrence of FMDV in livestock populations. We also show that the circulation of FMDV in buffalo is influenced by fine-scale geographic features, such as rivers, as well as herd-level factors. Our ability to discern these patterns is due to the large sample size of VP1 sequences recovered from this population. Thus, this project significantly advances knowledge about the ecology of FMDV in buffalo and the molecular and spatial epidemiology of FMDV in Eastern Africa.

## Acknowledgements

We are grateful to the National Academy of Sciences (NAS) and USAID for the funding entered through the Prime Agreement Number AID-OAA-A-11-00012. We are grateful for the management support from the Director, Kenya wildlife Service, FMD-Lab in Kenya, and Ol Pejeta Conservancy. We appreciate the technical field assistance from Joseph Kabugi, Elsie Maina, Dr. Stephen Ngulu, John Kariuki, Edward Kingori, and Stephanie Hauver Lemaiyan. This research was also supported in part by USDA- ARS CRIS 8064- 32000- 061- 00D.

